# seGMM: a new tool to infer sex from massively parallel sequencing data

**DOI:** 10.1101/2021.12.16.472877

**Authors:** Sihan Liu, Yuanyuan Zeng, Meilin Chen, Qian Zhang, Lanchen Wang, Chao Wang, Yu Lu, Hui Guo, Fengxiao Bu

## Abstract

Inspecting concordance between self-reported sex and genotype-inferred sex from genomic data is a significant quality control measure in clinical genetic testing. Numerous tools have been developed to infer sex for genotyping array, whole-exome sequencing, and whole-genome sequencing data. However, improvements in sex inference from targeted gene sequencing panels are warranted. Here, we propose a new tool, seGMM, which applies unsupervised clustering (Gaussian Mixture Model) to determine the gender of a sample from the called genotype data integrated aligned reads. seGMM consistently demonstrated > 99% sex inference accuracy in publicly available (1000 Genomes) and our in-house panel dataset, which achieved obviously better sex classification than existing popular tools. Compared to including features only in the X chromosome, our results show that adding additional features from Y chromosomes (e.g. reads mapped to the Y chromosome) can increase sex classification accuracy. Notably, for WES and WGS data, seGMM also has an extremely high degree of accuracy. Finally, we proved the ability of seGMM to infer sex in single patient or trio samples by combining with reference data and pinpointing potential sex chromosome abnormality samples. In general, seGMM provides a reproducible framework to infer sex from massively parallel sequencing data and has great promise in clinical genetics.

## 1 Introduction

Next-generation sequencing (NGS) has revolutionized the clinical field by transforming the landscape of clinical genetic testing and has been adopted as a standard for diagnosing hereditary disorders, expanding our understanding of clinical genetics, and offering new opportunities for personalized precision medicine over the last decade (Phillips and Douglas, 2018; Phillips et al., 2020). Clinical genetic testing usually refers to the analysis of DNA to identify pathogenic variants to aid in the diagnosis of disease (McPherson, 2006). It may focus on a single gene, multi-gene panels [targeted gene sequencing (TGS)], whole exome [whole exome sequencing (WES)], or whole genome [whole genome sequencing (WGS)](Di Resta et al., 2018). TGS is highly recommended in genetic testing because of its validity, utility, and cost-effectiveness, especially in hearing loss, cardiovascular disorders, and renal disorders (Lin et al., 2012; Saudi Mendeliome, 2015).

Parallelized TGS analysis of patients from the large cohort is commonplace in clinical genetic testing. Considerable efforts are required for quality control (QC) and preprocessing of these data before detecting pathogenic variants (Lee et al., 2017). Mismatched genders indicate potential sample swap, pollution, sex chromosome abnormalities, or sequencing error, which will substantially lead to erroneous conclusions and affect treatment decisions (Taylor et al., 2015; Webster et al., 2019). Thus, one essential QC step is verifying concordance between self-reported sex and genotype-inferred sex. Cytogenetic analyses, such as karyotyping, are gold standard methods of inferring sex but are time and effort consuming. Leveraging computational tools to infer genotypic sex from sequencing data of X and Y chromosomes is a convenient and powerful alternative strategy to verify sex concordance.

Several tools, such as PLINK, seXY, and XYalign, have been developed to infer sex using data from genotyping array, WES, or WGS. PLINK calculated the F coefficient with X chromosome heterozygosity to infer sex for genotype array data (Purcell et al., 2007). In contrast, seXY considered both X chromosome heterozygosity and Y chromosome missingness to infer sex in genotype array data by logistic regression (Qian et al., 2017). In particular, XYalign extract read count mapped to sex chromosomes and calculated the ratio of X and Y counts (Webster et al., 2019). Together with self-reported sex and calculated ratios, by plotting a scatter plot, users can infer sex with eyeballs for WES and WGS data. However, the accuracy of these methods in TGS panel data has not yet been fully evaluated, and improvements in sex inference from gene panel data are warranted.

In this study, we propose a new sex inference tool, seGMM, that determines the gender of a sample from called genotype data integrated aligned reads and jointly considers information on the X and Y chromosomes in diverse genomic data, including TGS panel data. seGMM applies Gaussian Mixture Model (GMM) clustering to classify the samples into different clusters. Compared to previous methods that use logistic regression and training data to infer sex, seGMM is more powerful for modeling data with different covariance structures and different numbers of mixture components for various genomic data.

## 2 Materials and Methods

### 2.1 Data

To evaluate the accuracy of existing methods and seGMM in inferring sex for TGS panel data, we used 2 datasets from publicly available sources (Supplementary Table S1) and 1 dataset from our in-house resource: (1) exon-targeted sequencing data for 1000 genes from 110 males and 98 females from the 1000 Genomes Project (Dataset 1, (Genomes Project et al., 2010)); (2) massively parallel sequencing of 785 deafness-related genes (Supplementary Table S2) from 8,950 males and 7,737 females (Dataset 2); and (3) targeted sequencing data for 189 autism risk genes across a cohort from the Autism Clinical and Genetic Resources in China (ACGC), including 42 females and 205 males (Dataset 3, (Guo et al., 2018)).

In addition, to assess the application of seGMM in inferring sex for WES and WGS data, we used 2 datasets from publicly available sources (Supplementary Table S3 and Supplementary Table S4) and 1 dataset from our in-house source: (1) exome sequencing data from 164 males and 118 females from the 1000 Genomes Project (Dataset 4, (Genomes Project et al., 2015)); (2) 27 high-coverage whole genomes from the 1000 Genomes Project including 11 males and 16 females (Dataset 5; (Genomes Project et al., 2015)) and (5) exome sequencing data from 1,255 males and 1,138 females from our in-house resource (Dataset 6).

The publicly available BAM files were previously mapped to the reference genome (GRCh37), which we can directly use for downstream analyses. For our in-house datasets, Fastp was used to remove adapters and low-quality reads, as well as evaluate the quality of sequencing data via several measures, including Q20, sequence duplication levels, coverage, and GC content (Chen et al., 2018). After evaluation, none of the samples were excluded. Clean DNA sequencing reads were mapped to the human reference genome hg19 using the BWA-MEM algorithm (Li and Durbin, 2009). Duplicated reads for public BAM files and our in-house BAM files were removed using PicardTools. Genomic variants were called following the Genome Analysis Toolkit software best practices (McKenna et al., 2010). Variants were filtered by VCFtools (Danecek et al., 2011) with (1) missing in more than 50% of samples; (2) minor allele count < 3; (3) quality < 30 and (4) DP < 5.

### 2.2 Inferring genetic sex with seGMM

As expected, five features may be associated with sex, including X chromosome heterozygosity (XH), reads mapped to the X chromosome (Xmap), reads mapped to the Y chromosome (Ymap), the ratio of X/Y counts (XYratio), and the mean depth of exons in the sex-determining region of the Y chromosome (SRY) gene (SRY_dep). Usually, seGMM computes XH as the fraction of all genotypes on the X chromosome with two different allele calls, excluding missing genotypes. Xmap/Ymap was computed as the fraction of high-quality reads (mapq>30) that mapped to the X/Y chromosome in all high-quality reads that mapped to the genome with samtools (Li et al., 2009). XYratio was computed as the ratio of Xmap divided by Ymap. SRY_dep was extracted by mosdepth with high-quality reads (mapq>30) (Pedersen and Quinlan, 2018). Considering that the panel data may only contain genes in the X or Y chromosome, seGMM allows users to select features put into the GMM model.

After extracting features from BAM and VCF files, features were normalized to the same level using the scale function in R. Then, the R package mclust was used to perform model-based clustering via the EM algorithm to classify the samples into different clusters (Scrucca et al., 2016). Samples with an uncertainty greater than 0.1 were considered outliers. In addition, seGMM can infer genetic sex for a single patient or trio samples with additional reference data containing the same features selected to put into the model (Figure 1).

**Fig. 1.**
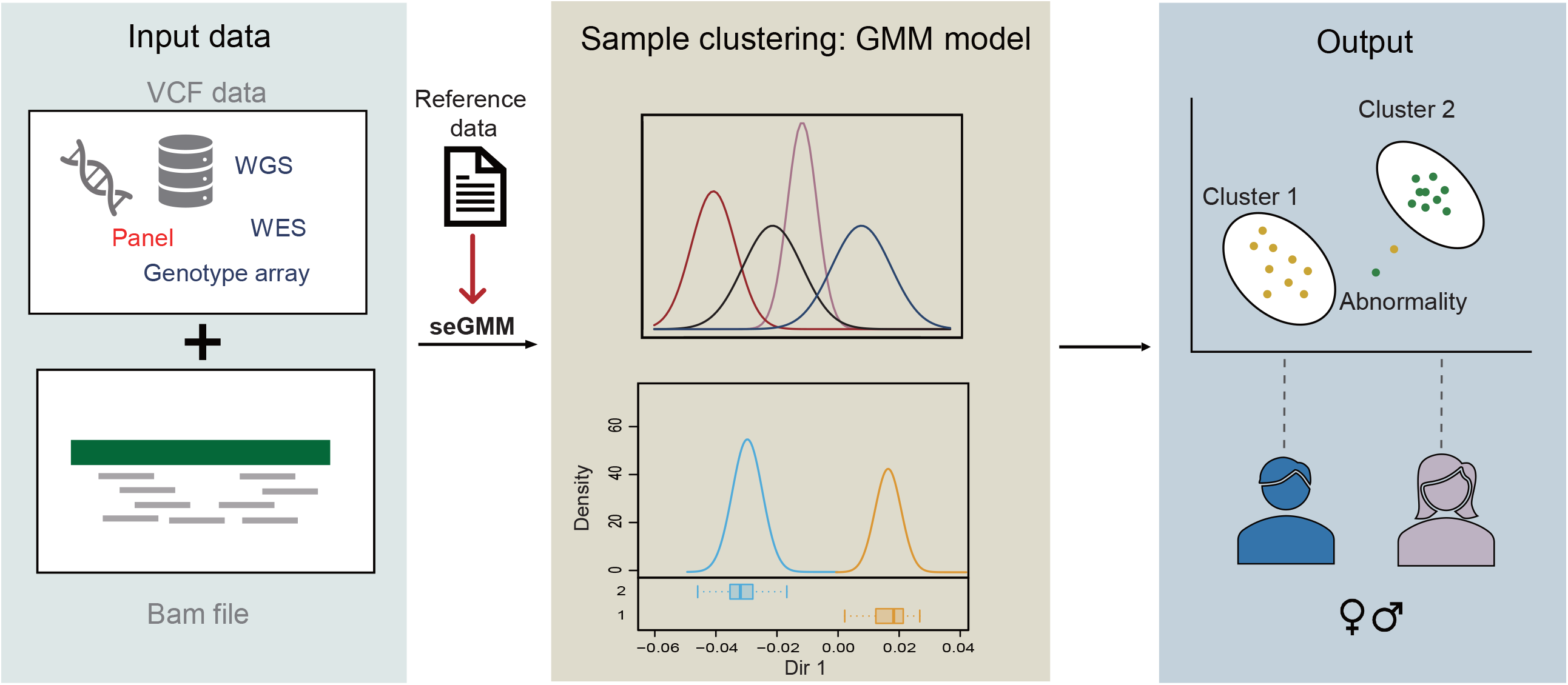
Schematic diagram of seGMM. The input file to seGMM is VCF and BAM file, seGMM will automatically collect features and build the GMM model.

### 2.3 Identifying potential sex abnormity samples

To pinpoint sex abnormity samples, we first define these values as mean_xmap_z, sd_xmap_z, mean_ymap_z and sd_ymap_z, where z □ {m, f} denotes whether the summaries were conditioned on the genetically determined males or females. Then, we defined the following six gates to classify individuals according to the values above:

- XY gate:
  - mean_xmap_m - 3 sd_xmap_m < x < mean_xmap_m + 3 sd_xmap_m
  - mean_ymap_m - 3 sd_ymap_m < y < mean_ymap_m + 3 sd_ymap_m

- XYY gate:
  - mean_xmap_m - 3 sd_xmap_m < x < mean_xmap_m + 3 sd_xmap_m
  - y > 2 mean_ymap_m

- XX gate:
  - mean_xmap_f - 3 sd_xmap_f < x < mean_xmap_f+ 3 sd_xmap_f
  - mean_ymap_f - 3 sd_ymap_f < y < mean_ymap_f + 3 sd_ymap_f

- XXY gate:
  - x > 2 mean_xmap_f
  - mean_ymap_m-3 sd_ymap_m < y < mean_ymap_m + 3 sd_ymap_m

- XXX gate:
  - x > 3 mean_xmap_f
  - mean_ymap_f - 3 sd_ymap_f < y < mean_ymap_f + 3 sd_ymap_f

- X gate:
  - x < 0.5 mean_xmap_f
  - mean_ymap_f - 3 sd_ymap_f < y < mean_ymap_f + 3 sd_ymap_f

### 2.4 Comparing performance with existing methods

To compare the performance between seGMM and existing methods. First, we downloaded and configured three tools for sex inference: PLINK 1.9, XYalign and seXY. For PLINK 1.9, X chromosome pseudoautosomal region was first splited off with --split-x. Then, --check-sex was running once without parameters, eyeball the distribution of F estimates, and rerun with parameters corresponding to the empirical gap. For XYalign, following the method described in their published paper, we used the CHROM_STATS module to obtain the depth of the 19 chromosome, X chromosome and Y chromosome. Then, the depth of the X and Y chromosomes was normalized by dividing it by the depth of chromosome 19. Finally, we plotted a scatter plot to assess sex-mismatched samples. For seXY, the first step was obtain the X.ped and Y.ped files with PLINK. Next, sex inference was conducted with seXY using X.ped, Y.ped and the training data set (subjects in prostate cancer and ovarian cancer GWAS) provided by seXY. PLINK was applied to all datasets. Since the target gene panel data of Datasets 2 and 3 do not contain genes located on the Y chromosome, XYalign and seXY were only applied to Dataset 1. In addition, XYalign was also applied to WES and WGS data.

### 2.5 STR analysis for verifying sex

The STR analysis was conducted in our own-designed multiplex STR system (modified based on PowerPlex® 16 System), which allows coamplification and four-color detection of sixteen loci (fifteen STR loci and Amelogenin). The primers for Amelogenin were designed as 5’-GTTCAGACGTGTGCTTCAACTTCAGCTATGAGGTAATTTTTC – 3’ and 5’-ATCCGACGGTAGTGTCCAACCATCAGAGCTTAAACTGG – 3’. All sixteen loci were amplified simultaneously in a single tube and analyzed in a single injection or gel lane. One primer for each of the vWA, amelogenin, FGA and TPOX loci was labeled with carboxyrhodamine (ROX). The amplicons were separated on an ABI 3730XL Genetic Analyzer, and data were collected using GeenMapper ID v3.2. Since females are XX, only a single peak is observed when testing female DNA, whereas males, which possess both X and Y chromosomes, exhibit two peaks with a 6 bp difference.

## 3 Results

### 3.1 seGMM achieved better sex classification in TGS data than existing methods

We calculated XH, Xmap, Ymap and XYratio for Dataset 1. We found that Ymap and XYratio plot as distinct clusters for the majority of males and females (Figure 2). The distribution of XH and Xmap for males and females has a lot of intersections, suggesting that the accuracy of sex inference is limited if we only include features extracted from the X chromosome. Our results proved this hypothesis: with features extracted only from the X chromosome, seGMM reported 59 samples as outliers, and the accuracy for the remaining samples was only 84.56%. Next, we evaluate the performance of seGMM in Dataset 1 using the 4 features we calculated before. We found that no samples were reported as outliers, and the accuracy increased to 99.52% after including features from the Y chromosome (Table 1). Moreover, looking into different genders, the accuracy of seGMM in females was 98.98%, and that in males was 100%. The only female sample (NA19054) misclassified by seGMM had an XY ratio that closely mirrored those of males.

**Table 1.** Accuracy of different methods in inferring sex with Dataset 1.

|  | Accuracy for all samples (%) | Accuracy for Male (%) | Accuracy for Female (%) |
| --- | --- | --- | --- |
| PLINK | 81.44 | 48.28 | 100 |
| seXY | 62.5 | 45.45 | 81.63 |
| seGMM | 99.52 | 100 | 98.98 |

**Fig. 2.**
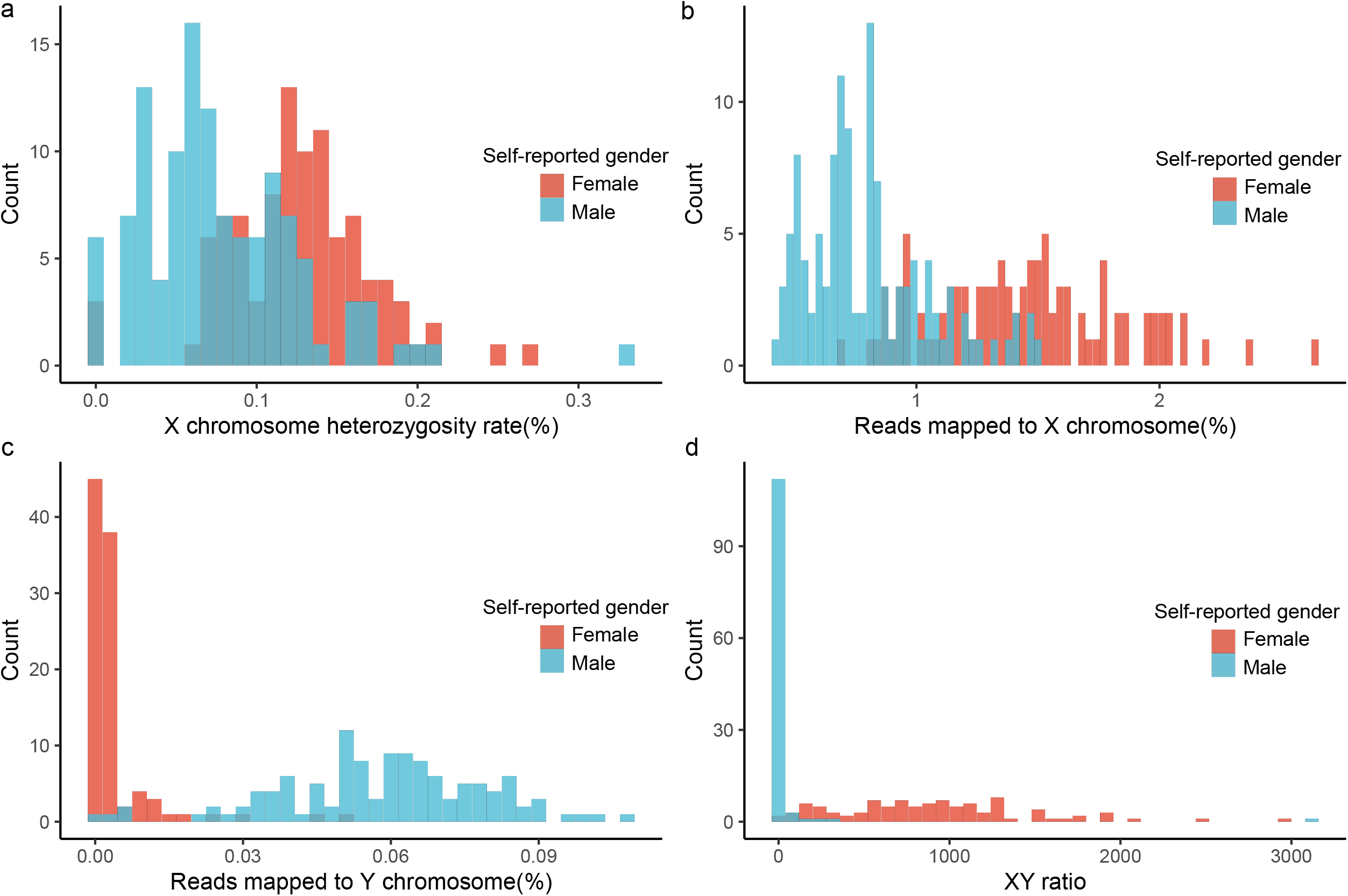
Distribution of features collected from Dataset 1. a. Distribution of X chromosome heterozygosity rate (%) between males and females. b. Distribution of reads mapped to the X chromosome (%) between males and females. c. Distribution of reads mapped to the Y chromosome (%) between males and females. d. Distribution of XYratio between male and female.

To assess the performance of seGMM, we also applied PLINK, seXY and XYalign in Dataset 1 to assess the accuracy of these existing tools in inferring sex. We discovered that the distribution of the F coefficient is concentrated between 0-0.9 and without an empirical gap (Supplementary Figure S1). The accuracy of PLINK was 81.44 %, with 94 samples whose predicted sex was clear (Table 1). The accuracy of seXY is only 62.5%. For XYalign, which does not directly provide a predicted sex, we plot the normalized sequence depth of chromosome X and chromosome Y. A couple of females and males are mixed, indicating the loss of accuracy with XYalign compared to seGMM (Supplementary Figure S2). Moreover, we have tested the computation time of different methods using 1 core, 10 cores and 20 cores on a server with 64 Intel(R) Xeon(R) CPU E7-8895 v3 @ 2.60 GHz. We can see that, as expected, seGMM costs much more time than PLINK and seXY, which don’t collect features of reads mapped to the X and Y chromosomes. Contrary to PLINK and seXY, with 1 core, seGMM costs fairly time compared to XYalign, while when using 20 cores, seGMM achieves 10 times faster than XYalign. (Supplementary Table S5)

To validate the performance of seGMM in inferring sex from gene panel data, we further applied seGMM to Dataset 2 and Dataset 3. Since the target gene panel data of these datasets do not contain genes located on the Y chromosome, we calculated XH and Xmap as features. In contrast to Dataset 1, the distribution of XH and Xmap for Dataset 2 plot as distinct clusters for the majority of males and females (Figure 3). We compared the performance of seGMM in inferring sex with PLINK. In total, the accuracy of seGMM was 99.10%, while the accuracy of PLINK was 86.58% (Table 2). Additionally, looking into different genders, the accuracy of seGMM in females was nearly 100% and 98.34% in males. The few males misclassified by seGMM have XH values that closely mirror those of females. Furthermore, the accuracy of seGMM in Dataset 3 is 92.31% (Table 2), while the accuracy of PLINK is relatively lower (38.87%).

**Table 2.** Accuracy of different methods in inferring sex with Dataset 2 and Dataset 3.

|  | Dataset 2 | Dataset 3 |
| --- | --- | --- |
| PLINK | 86.58 | 38.87 |
| seGMM | 99.10 | 92.31 |

**Fig. 3.**
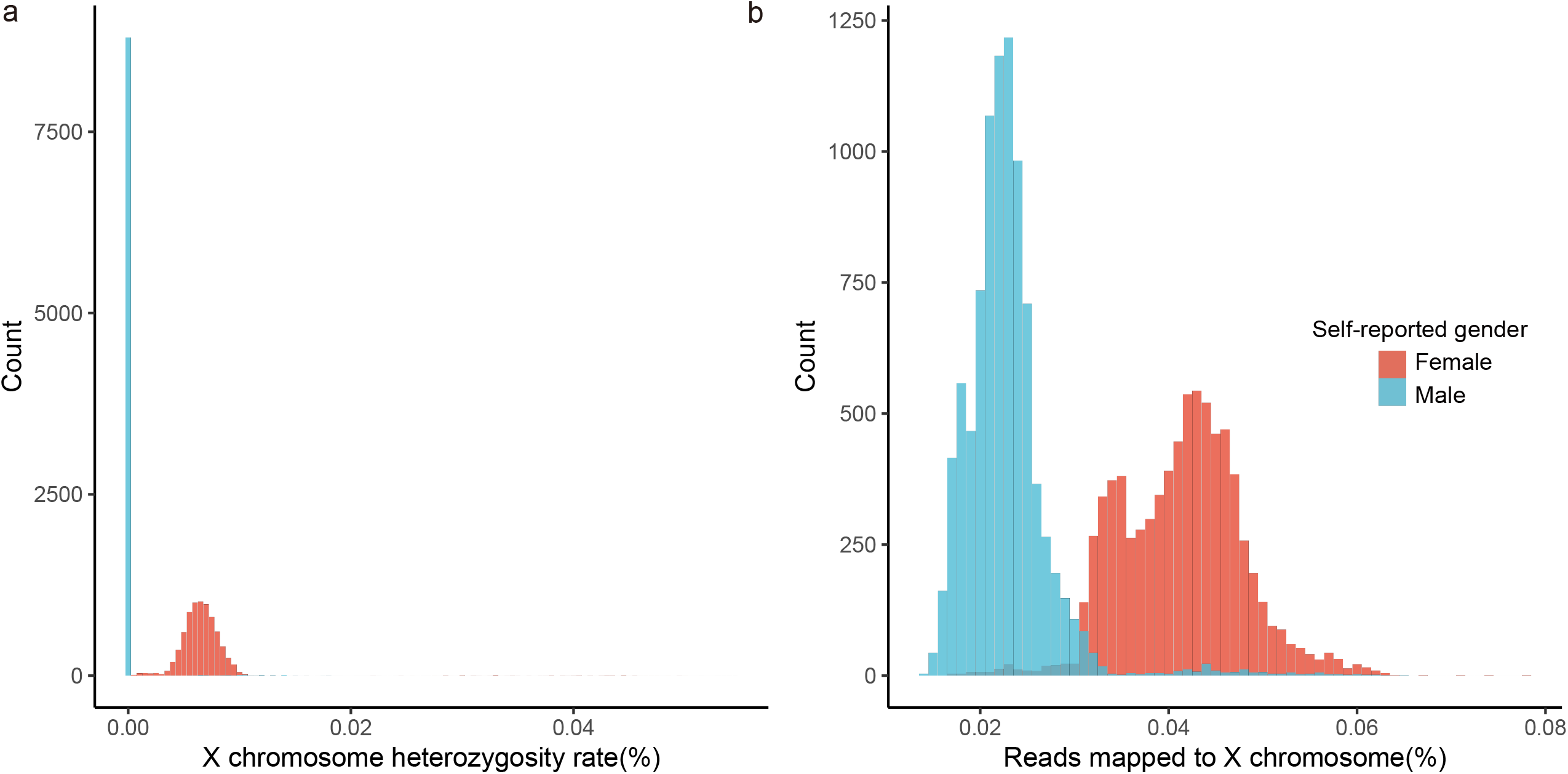
Distribution of features collected from Dataset 2. a. Distribution of X chromosome heterozygosity rate (%) between males and females. b. Distribution of reads mapped to the X chromosome (%) between males and females.

### 3.2 seGMM has good performance in inferring sex for WES and WGS data

Recently, WES and WGS have shown promise in becoming a first-tier diagnostic test for patients with Mendelian disorders. Therefore, we evaluated the performance of seGMM in inferring sex from WES and WGS data. First, we applied seGMM to publicly available WES data (Dataset 4). The accuracy of seGMM was 100%, indicating that seGMM also has excellent performance in WES data (Table 3 and Supplementary Figure S3). Meanwhile, the accuracy of PLINK was 100%, while the accuracy of XYalign was 99.65%. NA12413 was mixed with female samples (Supplementary Figure S4).

**Table 3.** Accuracy of different methods in inferring sex with WES and WGS data.

|  | PLINK | XYalign | seGMM |
| --- | --- | --- | --- |
| 1000G phase3 WES data | 100 | 99.65 | 100 |
| 1000G phase3 high quality WGS data | 100 | 100 | 100 |
| In-house WES data | 99.54 | 99.66 | 99.75 |

In addition, we applied seGMM to our in-house WES data (Dataset 6). The accuracy of seGMM in our in-house WES data was 99.75%, 99.76% for males and 99.74% for females (Supplementary Table S6). Six samples (3 males and 3 females; Table 3 and Figure 4a) were mismatched between SNP-inferred sex and self-reported sex, indicating potential misregistration of clinical information. For PLINK, the accuracy was 99.54%, and 11 samples were mismatched (Table 3 and Figure 4b). The accuracy for XYalign was 99.66%, and 8 samples were mismatched (Figure 4c). Six samples were detected with mismatched sex in the three methods; however, 5 mismatched samples detected with PLINK and 2 mismatched samples detected with XYalign were correct in seGMM (Figure 4d).

**Fig. 4.**
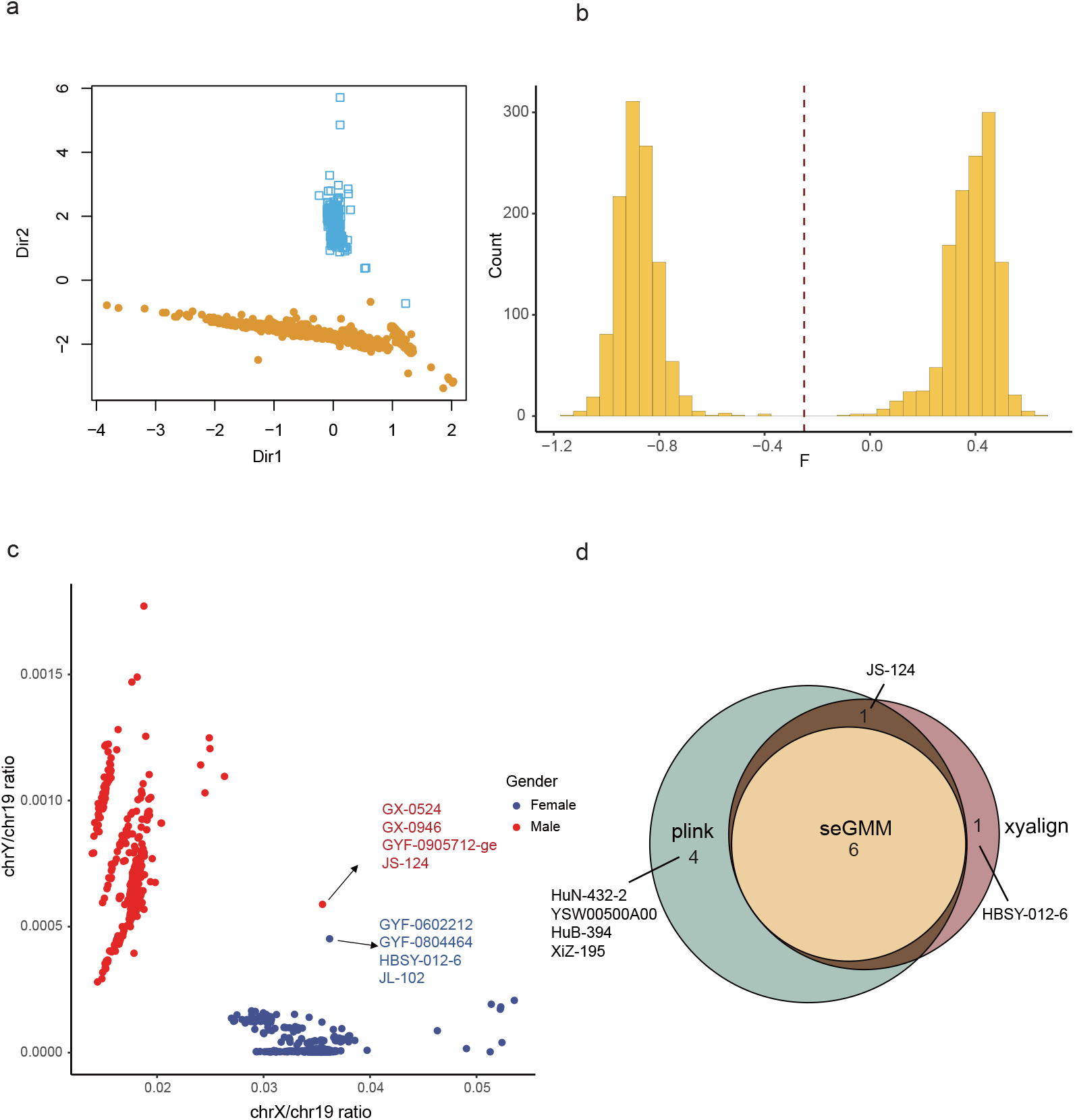
Accuracy of different methods in inferring sex with our in-house WES data. a. Sample clustering results of seGMM. b. Distribution of F coefficient. c. Scatter plot of normalized X and Y ratio using XYalign. d. Venn plot of mismatched gender samples detected by the three methods.

To verify the real sex of these 6 samples, we performed STR analysis with a sex marker, the amelogenin gene. PCR products generated from the amelogenin gene are widely accepted for use in sex identification. The amelogenin gene is highly conserved and occurs on both the X- and Y-chromosomes. With a 6 bp deletion of the amelogenin gene in the Y chromosome, amplicons generated from the X and Y chromosomes were distinguished from one another when electrophoretic separation was performed to separate STR alleles. The results showed that the real sex matched the seGMM prediction results, proving the pinpoint accuracy of seGMM (Table 4 and Figure 5). Finally, seGMM was conducted on the WGS data (Dataset 5), and the accuracy was 100% (Table 3). The accuracy for PLINK and XYalign is also 100%.

**Table 4.** Accuracy of different methods in inferring sex with WES and WGS data.

| Sample ID | Amelogenin | Self-reported gender | seGMM inferred gender | Experiment validate gender |
| --- | --- | --- | --- | --- |
| GX-0524 | 209.15 | Male | Female | Female |
|  | - |  |  |  |
| GX-0946 | 209.06 | Male | Female | Female |
|  | - |  |  |  |
| GYF-0602212 | 209.04 | Female | Male | Male |
|  | 214.8 |  |  |  |
| GYF-0804464 | 209.06 | Female | Male | Male |
|  | 214.85 |  |  |  |
| GYF-0905712-ge | 209.11 | Male | Female | Female |
|  | - |  |  |  |
| JL-102 | 209.18 | Female | Male | Male |
|  | 214.92 |  |  |  |

**Fig. 5.**
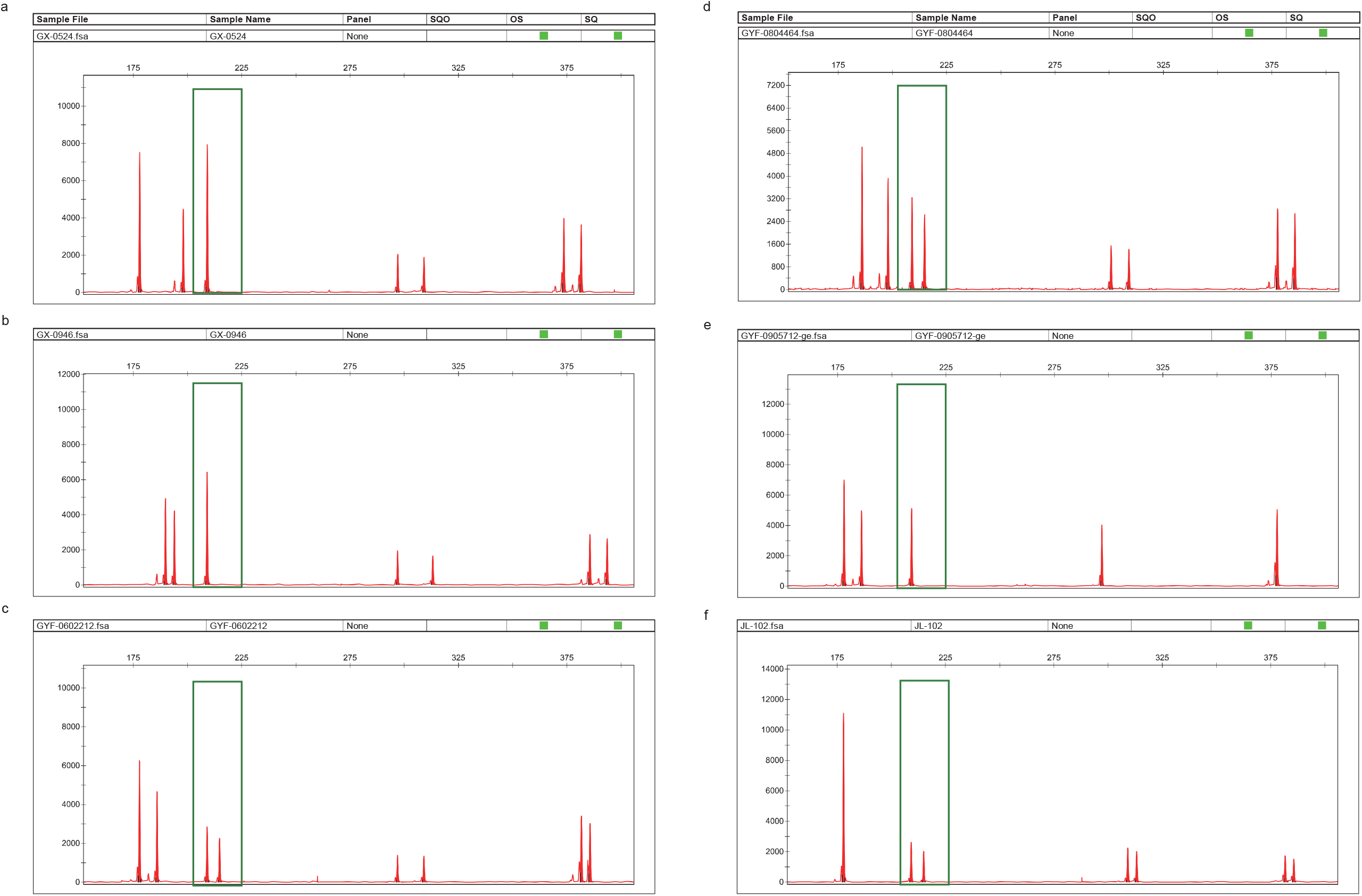
Experimentally verified gender results.

### 3.3 The ability of seGMM in clinical application

In clinical practice, individual patient or trio samples are usually sequenced to obtain a molecular diagnosis. However, the GMM model requires a sufficient sample size to ensure the accuracy of classification. To address this problem, seGMM permits users to provide additional reference data. By combining the features from reference data, seGMM can ensure accuracy for clinical application. Taking WES data as an example, using features including Xmap, Ymap, XYratio and normalized SRY_dep (divided by XYratio), we found that with 1000 Genomes data points as a reference, all samples in our in-house WES data were predicted accurately and vice versa.

Additionally, approximately 0.25% male and 0.15% female live births demonstrated some form of sex chromosome abnormality. A previous study examined the feasibility of defining five gates to classify individuals according to the normalized X and Y chromosome ratio, calculated on genetically determined males and females, respectively (Turro et al., 2020). Similarly, following this strategy, seGMM can automatically classify samples into 6 sex chromosome karyotypes (XX, XY, XYY, XXY, XXX and X) according to the Xmap and Y map. For publicly available WES and WGS data, none of the samples had sex chromosome abnormalities. For our in-house WES data, 3 samples (HBSY-012-ge, HBSY-012 and XiZ-086) were identified with XYY chromosome karyotypes.

## 4 Discussion

This article introduces a sex inference tool, seGMM, to infer genotype sex from NGS data, especially TGS panel data. seGMM integrates several tools and algorithms into a single workflow. Using features extracted from sex chromosomes, seGMM applied unsupervised clustering to classify the samples. Compared to many existing supervised methods that attempt to infer sex by training a logistic regression classifier based on limited available data, seGMM can be applied directly to different types of genomic data. By comparing PLINK, seXY and XYalign, we proved that seGMM outperforms existing tools in inferring sex with TGS panel data and has excellent performance in WES and WGS data.

Additionally, compared with including features extracted from only the X chromosome, we discovered that jointing valuable information on the Y chromosome improved the accuracy of inferring sex. Our data suggest that adding probes targeting unique regions of the Y chromosome, particularly the exon of the SRY, which is involved in male-typical sex development (Gubbay et al., 1990; Parma and Radi, 2012), is helpful in inferring genders using TGS panel data.

One important innovation of seGMM is that seGMM is adapted to clinical applications that can be applied to individual patients and automatically report sex chromosome abnormality samples. When applying seGMM with reference data, attention should be given to the consistency of the analytical methods between reference data and testing data, since inconsistent analysis methods can introduce bias.

There are several limitations for seGMM. First, as a method expected to find sex chromosome abnormal samples, seGMM classifies the samples by calculating the Xmap and Ymap intervals in which they are located. However, the interval is fixed and when the number of male or female samples in the testing dataset is small, the distribution of the Xmap/Ymap is approximately nonnormal and then the sex chromosome karyotypes classification of samples in this case may be inaccurate. Hence, we recommended that if the sample size of the male or female is small, combining with a reference data is an effective strategy to ensure the accuracy of results. Second, the computational time of seGMM depends on many factors, such as the number of features, the number of samples, and the number of threads used. seGMM costs much time in collecting features of reads mapped to the X and Y chromosomes, causing it to run slower than PLINK and seXY but faster than XYalign. The running speed of seGMM can increased by adding cores, and we also consider rewriting it as a GPU program to speed up in the future.

In conclusion, we have developed a new tool to infer genetic sex based on a Gaussian Mixture Model called seGMM, which combines stable predictive ability and clinical application. In addition, when the genomic data are TGS, seGMM is one of the best choices for inferring sex, which could meet the needs of clinical genetics.

## 5 Data Availability Statement

seGMM is publicly available at https://github.com/liusihan/seGMM and can be installed directly by Conda. Users can also download the source code from GitHub or PyPI and install related software. The data used in this study were retrieved from the 1000 Genomes database (https://ftp-trace.ncbi.nih.gov/1000genomes/ftp/). Further inquiries can be directed to the corresponding authors.

## 7 Author Contributions

SL developed the tool and drafted the manuscript. YZ, MC and QZ participated in data preprocessing and testing. LW and CW performed the experiment. YL and HG reviewed and revised the manuscript. FB designed and supervised the study and reviewed the manuscript. All authors contributed to the article and approved the submitted version.

