## Supplementary Material for "seGMM: a new tool to infer sex from massively parallel sequencing data"

### 1 Supplementary Figures and Tables

#### 1.1 Supplementary Figures

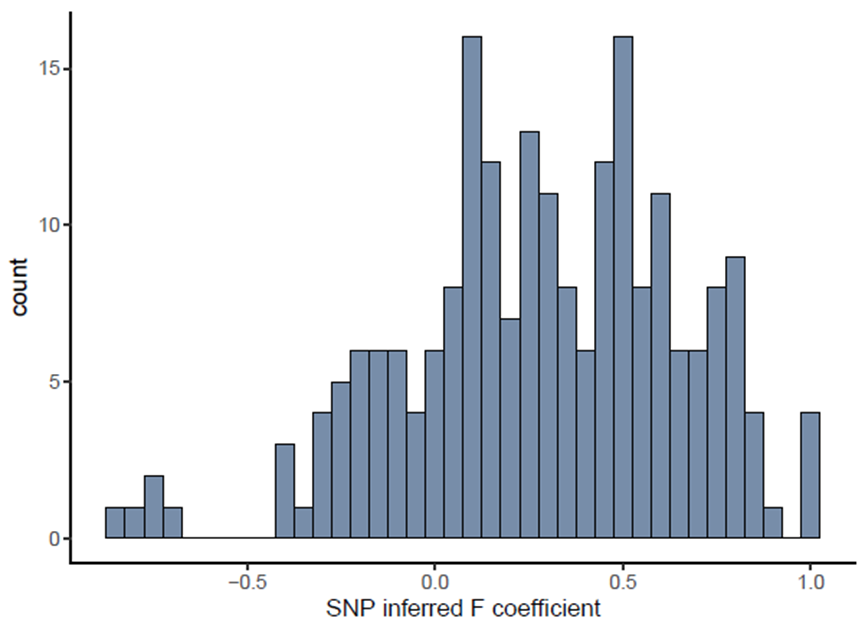

**Supplementary Figure 1.** Distribution of SNP inferred F coefficient collected from dataset 1 using plink.

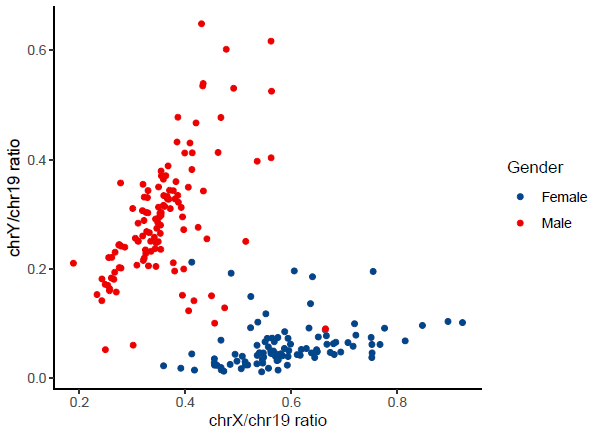

**Supplementary Figure 2.** Scatter plot of normalized X and Y ratio collected from dataset 1 using xyalign.

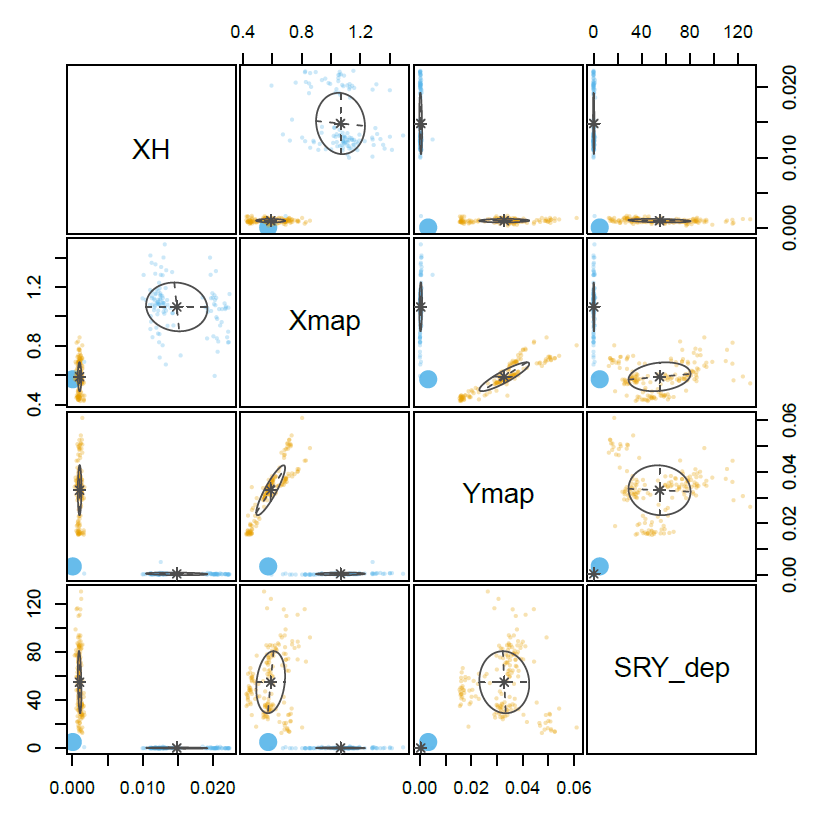

**Supplementary Figure 3.** Clustering of samples from Dataset 4 using seGMM.

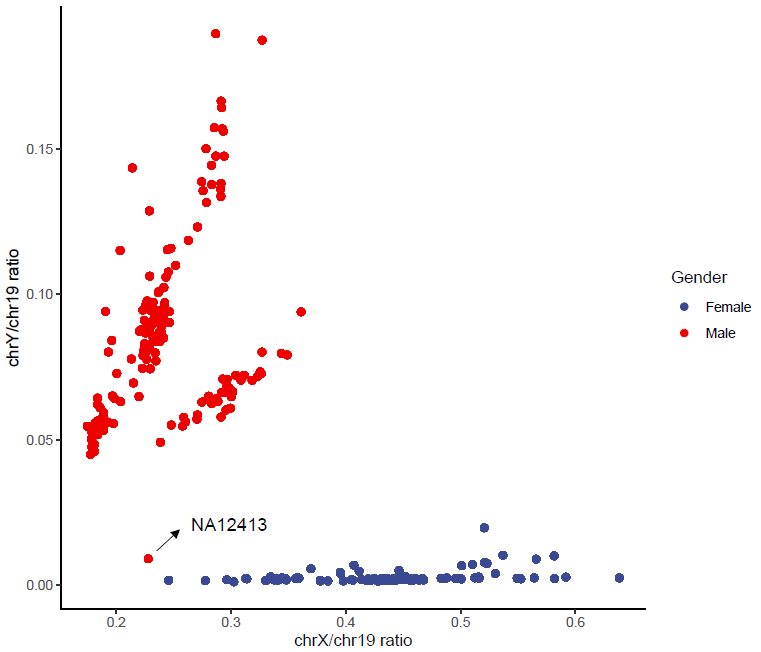

**Supplementary Figure 4.** Scatter plot of normalized X and Y ratio collected from dataset 4 using xyalign.

#### 1.2 Supplementary tables

**Supplementary** **Table 1.** Sex information for Dataset 1

| NA07000 | Female | NA07037 | Female | NA07048 | Male | NA07051 | Male |
| --- | --- | --- | --- | --- | --- | --- | --- |
| NA07347 | Male | NA07357 | Male | NA10847 | Female | NA10851 | Male |
| NA11830 | Female | NA11843 | Male | NA11893 | Male | NA11918 | Female |
| NA11919 | Male | NA11930 | Male | NA12005 | Male | NA12043 | Male |
| NA12144 | Male | NA12154 | Male | NA12234 | Female | NA12249 | Female |
| NA12272 | Male | NA12273 | Female | NA12275 | Female | NA12283 | Female |
| NA12286 | Male | NA12287 | Female | NA12347 | Male | NA12348 | Female |
| NA12383 | Female | NA12546 | Male | NA12716 | Male | NA12749 | Female |
| NA12775 | Male | NA12776 | Female | NA12827 | Male | NA12828 | Female |
| NA12830 | Female | NA12878 | Female | NA12890 | Female | NA18113 | Male |
| NA18126 | Female | NA18145 | Male | NA18164 | Female | NA18486 | Male |
| NA18487 | Male | NA18488 | Female | NA18498 | Male | NA18505 | Female |
| NA18508 | Female | NA18510 | Male | NA18519 | Male | NA18520 | Female |
| NA18522 | Male | NA18523 | Female | NA18526 | Female | NA18532 | Female |
| NA18537 | Female | NA18547 | Female | NA18552 | Female | NA18561 | Male |
| NA18563 | Male | NA18566 | Female | NA18582 | Female | NA18603 | Male |
| NA18608 | Male | NA18632 | Male | NA18633 | Male | NA18635 | Male |
| NA18636 | Male | NA18637 | Male | NA18638 | Male | NA18640 | Female |
| NA18641 | Female | NA18642 | Female | NA18643 | Male | NA18645 | Male |
| NA18647 | Male | NA18669 | Male | NA18670 | Female | NA18671 | Female |
| NA18674 | Male | NA18679 | Male | NA18683 | Female | NA18685 | Male |
| NA18687 | Female | NA18689 | Male | NA18694 | Female | NA18696 | Female |
| NA18698 | Male | NA18699 | Female | NA18701 | Male | NA18704 | Female |
| NA18707 | Male | NA18708 | Male | NA18740 | Male | NA18745 | Male |
| NA18747 | Male | NA18748 | Male | NA18757 | Male | NA18853 | Male |
| NA18858 | Female | NA18861 | Female | NA18867 | Female | NA18868 | Male |
| NA18870 | Female | NA18871 | Male | NA18873 | Female | NA18874 | Male |
| NA18907 | Female | NA18908 | Male | NA18909 | Female | NA18910 | Male |
| NA18912 | Female | NA18916 | Female | NA18943 | Male | NA18948 | Male |
| NA18950 | Female | NA18951 | Female | NA18952 | Male | NA18960 | Male |
| NA18961 | Male | NA18964 | Female | NA18965 | Male | NA18973 | Female |
| NA18980 | Female | NA18982 | Male | NA18983 | Male | NA18984 | Male |
| NA18985 | Male | NA18986 | Male | NA18988 | Male | NA18989 | Male |
| NA18999 | Female | NA19000 | Male | NA19003 | Female | NA19006 | Male |
| NA19007 | Male | NA19011 | Female | NA19012 | Male | NA19054 | Female |
| NA19055 | Male | NA19056 | Male | NA19057 | Female | NA19058 | Male |
| NA19059 | Female | NA19060 | Male | NA19062 | Male | NA19063 | Male |
| NA19064 | Female | NA19065 | Female | NA19066 | Male | NA19067 | Male |
| NA19070 | Male | NA19087 | Female | NA19088 | Male | NA19089 | Male |
| NA19091 | Male | NA19092 | Male | NA19102 | Female | NA19108 | Female |
| NA19130 | Male | NA19189 | Male | NA19190 | Female | NA19213 | Male |
| NA19222 | Female | NA19225 | Female | NA19235 | Female | NA19236 | Male |
| NA19238 | Female | NA19239 | Male | NA19240 | Female | NA19247 | Female |
| NA19248 | Male | NA19257 | Female | NA19546 | Female | NA19550 | Female |
| NA19551 | Female | NA19552 | Female | NA19553 | Female | NA19554 | Female |
| NA19555 | Male | NA19556 | Male | NA19558 | Male | NA19559 | Male |
| NA19560 | Female | NA19561 | Female | NA19562 | Female | NA19563 | Female |
| NA19564 | Female | NA19565 | Female | NA19566 | Female | NA19568 | Female |
| NA19569 | Female | NA19572 | Female | NA19573 | Female | NA19574 | Female |
| NA20504 | Female | NA20509 | Male | NA20510 | Male | NA20511 | Male |
| NA20513 | Male | NA20515 | Male | NA20516 | Male | NA20517 | Female |
| NA20520 | Male | NA20521 | Male | NA20522 | Female | NA20525 | Male |

**Supplementary** **Table 2.** Sample information for our in-house datasets.

|  | Male | Female | Total |
| --- | --- | --- | --- |
| Panel data | 8950 | 7737 | 16687 |
| WES | 1257 | 1136 | 2393 |

**Supplementary** **Table 3**. Sex information for Dataset 4

| NA06986 | Male | NA07000 | Female | NA07037 | Female |
| --- | --- | --- | --- | --- | --- |
| NA07048 | Male | NA07051 | Male | NA07347 | Male |
| NA07357 | Male | NA10847 | Female | NA10851 | Male |
| NA11829 | Male | NA11830 | Female | NA11831 | Male |
| NA11832 | Female | NA11843 | Male | NA11893 | Male |
| NA11918 | Female | NA11919 | Male | NA11920 | Female |
| NA11930 | Male | NA11992 | Male | NA11994 | Male |
| NA12003 | Male | NA12005 | Male | NA12043 | Male |
| NA12045 | Male | NA12144 | Male | NA12154 | Male |
| NA12155 | Male | NA12234 | Female | NA12249 | Female |
| NA12272 | Male | NA12273 | Female | NA12275 | Female |
| NA12282 | Male | NA12283 | Female | NA12286 | Male |
| NA12287 | Female | NA12347 | Male | NA12348 | Female |
| NA12383 | Female | NA12399 | Male | NA12400 | Female |
| NA12413 | Male | NA12546 | Male | NA12716 | Male |
| NA12717 | Female | NA12718 | Female | NA12748 | Male |
| NA12749 | Female | NA12750 | Male | NA12751 | Female |
| NA12761 | Female | NA12763 | Female | NA12775 | Male |
| NA12776 | Female | NA12827 | Male | NA12828 | Female |
| NA12829 | Male | NA12830 | Female | NA12842 | Male |
| NA12878 | Female | NA12889 | Male | NA12890 | Female |
| NA18486 | Male | NA18488 | Female | NA18489 | Female |
| NA18498 | Male | NA18499 | Female | NA18501 | Male |
| NA18504 | Male | NA18505 | Female | NA18508 | Female |
| NA18510 | Male | NA18516 | Male | NA18519 | Male |
| NA18520 | Female | NA18522 | Male | NA18523 | Female |
| NA18526 | Female | NA18530 | Male | NA18532 | Female |
| NA18534 | Male | NA18536 | Male | NA18537 | Female |
| NA18543 | Male | NA18544 | Male | NA18546 | Male |
| NA18547 | Female | NA18548 | Male | NA18549 | Male |
| NA18552 | Female | NA18557 | Male | NA18559 | Male |
| NA18561 | Male | NA18563 | Male | NA18566 | Female |
| NA18577 | Female | NA18579 | Female | NA18582 | Female |
| NA18595 | Female | NA18596 | Female | NA18597 | Female |
| NA18599 | Female | NA18602 | Female | NA18603 | Male |
| NA18608 | Male | NA18611 | Male | NA18612 | Male |
| NA18632 | Male | NA18633 | Male | NA18635 | Male |
| NA18636 | Male | NA18637 | Male | NA18638 | Male |
| NA18640 | Female | NA18641 | Female | NA18642 | Female |
| NA18643 | Male | NA18645 | Male | NA18647 | Male |
| NA18740 | Male | NA18745 | Male | NA18747 | Male |
| NA18748 | Male | NA18749 | Male | NA18757 | Male |
| NA18853 | Male | NA18856 | Male | NA18858 | Female |
| NA18861 | Female | NA18867 | Female | NA18868 | Male |
| NA18870 | Female | NA18871 | Male | NA18873 | Female |
| NA18874 | Male | NA18907 | Female | NA18908 | Male |
| NA18910 | Male | NA18912 | Female | NA18916 | Female |
| NA18917 | Male | NA18923 | Male | NA18924 | Female |
| NA18933 | Female | NA18934 | Male | NA18943 | Male |
| NA18948 | Male | NA18950 | Female | NA18951 | Female |
| NA18952 | Male | NA18953 | Male | NA18959 | Male |
| NA18960 | Male | NA18961 | Male | NA18964 | Female |
| NA18965 | Male | NA18966 | Male | NA18967 | Male |
| NA18968 | Female | NA18969 | Female | NA18970 | Male |
| NA18971 | Male | NA18972 | Female | NA18973 | Female |
| NA18974 | Male | NA18975 | Female | NA18976 | Female |
| NA18978 | Female | NA18980 | Female | NA18981 | Female |
| NA18982 | Male | NA18983 | Male | NA18984 | Male |
| NA18985 | Male | NA18986 | Male | NA18987 | Female |
| NA18988 | Male | NA18989 | Male | NA18990 | Male |
| NA18991 | Female | NA18999 | Female | NA19000 | Male |
| NA19003 | Female | NA19006 | Male | NA19007 | Male |
| NA19011 | Female | NA19012 | Male | NA19054 | Female |
| NA19055 | Male | NA19056 | Male | NA19057 | Female |
| NA19058 | Male | NA19059 | Female | NA19060 | Male |
| NA19062 | Male | NA19063 | Male | NA19064 | Female |
| NA19065 | Female | NA19066 | Male | NA19067 | Male |
| NA19070 | Male | NA19072 | Male | NA19074 | Female |
| NA19075 | Male | NA19076 | Male | NA19077 | Female |
| NA19078 | Female | NA19079 | Male | NA19080 | Female |
| NA19081 | Female | NA19082 | Male | NA19083 | Male |
| NA19084 | Female | NA19085 | Male | NA19086 | Male |
| NA19087 | Female | NA19088 | Male | NA19089 | Male |
| NA19091 | Male | NA19092 | Male | NA19098 | Male |
| NA19102 | Female | NA19108 | Female | NA19116 | Female |
| NA19119 | Male | NA19130 | Male | NA19131 | Female |
| NA19137 | Female | NA19138 | Male | NA19141 | Male |
| NA19143 | Female | NA19144 | Male | NA19152 | Female |
| NA19153 | Male | NA19159 | Female | NA19160 | Male |
| NA19171 | Male | NA19172 | Female | NA19189 | Male |
| NA19190 | Female | NA19197 | Female | NA19198 | Male |
| NA19200 | Male | NA19201 | Female | NA19204 | Female |
| NA19206 | Female | NA19207 | Male | NA19209 | Female |
| NA19210 | Male | NA19213 | Male | NA19222 | Female |
| NA19223 | Male | NA19225 | Female | NA19235 | Female |
| NA19236 | Male | NA19238 | Female | NA19239 | Male |
| NA19247 | Female | NA19248 | Male | NA19257 | Female |
| NA20504 | Female | NA20506 | Female | NA20508 | Female |
| NA20509 | Male | NA20510 | Male | NA20511 | Male |
| NA20512 | Male | NA20513 | Male | NA20515 | Male |
| NA20516 | Male | NA20517 | Female | NA20518 | Male |
| NA20519 | Male | NA20520 | Male | NA20521 | Male |
| NA20522 | Female | NA20524 | Male | NA20525 | Male |
| NA20527 | Male | NA20528 | Male | NA20529 | Female |

**Supplementary** **Table 4**. Sex information for Dataset 5

| HG00096 | Male | HG00268 | Female | HG00419 | Female |
| --- | --- | --- | --- | --- | --- |
| HG00759 | Female | HG01051 | Male | HG01112 | Male |
| HG01500 | Male | HG01565 | Male | HG01583 | Male |
| HG01595 | Female | HG01879 | Male | HG02568 | Female |
| HG02922 | Female | HG03006 | Male | HG03052 | Female |
| HG03642 | Female | HG03742 | Male | NA12878 | Female |
| NA18525 | Female | NA18939 | Female | NA19017 | Female |

**Supplementary** **Table 5**. Computation time of different methods in inferring sex with Dataset 1.

|  | 1 core | 10 cores | 20 cores |
| --- | --- | --- | --- |
| PLINK | ~3s | — | — |
| seXY | ~20s | — | — |
| XYalign | ~12.15m | ~11.8m | ~11.6m |
| seGMM | ~11m | ~8m | ~1m |

**Supplementary** **Table 6**. Accuracy of inferring sex for WES and WGS data using seGMM in different gender samples.

|  | All samples (%) | Male (%) | Female (%) |
| --- | --- | --- | --- |
| WES (1000G) | 100 | 100 | 100 |
| WGS (1000G) | 100 | 100 | 100 |
| In-house WES | 99.75 | 99.76 | 99.74 |
